# Perisaccadic perceptual mislocalization strength depends on the visual appearance of saccade targets

**DOI:** 10.1101/2024.08.16.608354

**Authors:** Matthias P. Baumann, Anna F. Denninger, Ziad M. Hafed

**Affiliations:** Werner Reichardt Centre for Integrative Neuroscience, University of Tübingen, Tübingen, Germany; Hertie Institute for Clinical Brain Research, University of Tübingen, Tübingen, Germany; Department for Psychiatry and Psychotherapy, University of Tübingen, Tübingen, Germany; Center for Mental Health (TüCMH), University of Tübingen, Tübingen, Germany

## Abstract

We normally perceive a stable visual environment despite repetitive eye movements. To achieve such stability, visual processing integrates information across saccades, and laboratory hallmarks of such integration are robustly observed by presenting brief perimovement visual probes. In one classic phenomenon, perceived probe locations are grossly erroneous. This phenomenon is believed to depend, at least in part, on corollary discharge associated with saccade-related neuronal movement commands. However, we recently found that superior colliculus motor bursts, a known source of corollary discharge, can be different for different image appearances of the saccade target. Therefore, here we investigated whether perisaccadic perceptual mislocalization also depends on saccade-target appearance. We asked human participants to generate saccades to either low (0.5 cycles/deg) or high (5 cycles/deg) spatial frequency gratings. We always placed a high contrast target spot at grating center, to ensure matched saccades across image types. We presented brief perisaccadic probes, which were high in contrast to avoid saccadic suppression, and the subjects pointed (via mouse cursor) at their perceived locations. We observed stronger perisaccadic mislocalization for low spatial frequency saccade targets, and for upper visual field probe locations. This was despite matched saccade metrics and kinematics across conditions, and it was also despite matched probe visibility for the different saccade target images (low versus high spatial frequency gratings). To the extent that perisaccadic perceptual mislocalization depends on corollary discharge, our results suggest that such discharge might relay more than just spatial saccade vectors to the visual system; saccade-target visual features can also be transmitted.

**Significance:** Brief visual probes are grossly mislocalized when presented in the temporal vicinity of saccades. While the mechanisms of such mislocalization are still under investigation, one component of them could derive from corollary discharge signals associated with saccade movement commands. Here, we were motivated by the observation that superior colliculus movement bursts, one source of corollary discharge, vary with saccade-target image appearance. If so, then perisaccadic mislocalization should also do so, which we confirmed.

## Introduction

Vision is a highly active process, continuously utilizing eye movements to both sample and modulate incoming retinal image streams. Because eye movements, such as saccades, necessarily shift retinal images even with a completely stable environment, perceptual processes bridging inter- and intra-saccadic image sequences are known to take place (1–4). One classic approach to the study of perisaccadic vision has been to present very brief visual probes before, during, and after saccades (5–11). Such presentation has led to two classic observations. First, there is a strong reduction in visual sensitivity to brief perisaccadic probes (7–9, 12–14), called perceptual saccadic suppression. Second, with high contrast probes, these probes are perceptually detected but grossly mislocalized in space (5, 6, 10, 11, 15–18), in a phenomenon called perisaccadic perceptual mislocalization.

Perisaccadic perceptual mislocalization has different properties depending on the visual and motor conditions under which it is observed. For example, the presence of visual reference frames makes the mislocalization appear like a compression towards the saccade target (15). That is, brief probes presented ahead of the saccade target are misperceived backwards in position, and probes presented nearer to the initial fixation position than the saccade target are misperceived forward in position (11). On the other hand, mislocalization is uniform in direction in complete darkness (5, 6, 15). In addition, with compression (which we focus on here), the two-dimensional landscape of perisaccadic perceptual mislocalization strongly depends on saccade direction, with upward saccades causing particularly large backwards compression for probes presented ahead of the saccade target (17).

Because the properties of perisaccadic perceptual mislocalization depend on the visual conditions, one neuronal mechanism underlying this phenomenon could be sensory in nature. For example, in some models, classic perisaccadic compression of space (11) can be easily explained using saccade-induced retinotopic shifts of probe representations from the periphery to the fovea; when these shifts occur on anatomically distorted topographic visual maps having foveal magnification, then compression is a simple consequence of readout from such anatomically distorted maps (19). Indeed, a simple modification of such models to include both foveal as well as upper visual field magnification can also conceptually account for the greater compression strength observed for upward saccades (17). Specifically, in a sensory-motor area such as the superior colliculus (SC), there exist both foveal (20) as well as upper visual field (21) magnification. Thus, if mislocalization depends on how retinotopic representations of probes are translated from the periphery to the fovea on anatomically distorted maps, then larger perisaccadic mislocalization for upward saccades would be expected if such maps additionally differentially represent the upper visual field (17).

Having said that, it is also generally assumed that perisaccadic perceptual mislocalization does have a motor component associated with it. Besides the different representations for upward saccades in the SC (21, 22), and the resultant implications on mislocalization strength (17), the SC is a known source of saccade-related corollary discharge signals (23–29). Such corollary discharge signals are sufficient to cause perisaccadic retinotopic remapping of visual response fields in the cortex (30). Thus, if such cortical remapping is indeed one neuronal basis for perceptual mislocalization (31, 32), despite some controversy (33–36), then studying the roles of SC saccade-related discharge in perceptual mislocalization is very worthwhile.

One interesting recently reported property of SC motor discharge is related to the saccade target’s visual appearance. Specifically, we recently observed that SC motor bursts can be different for different saccade target images, even under matched saccade metrics and kinematics (37). Since SC motor bursts are relayed to the cortex virtually unchanged (24), this implies that perisaccadic corollary discharge from the SC (23, 25, 30) may potentially relay not only the movement vector information to the cortex, but also integrated information about the visual appearance of the saccade target. If so, then perceptual phenomena that are believed to depend on perisaccadic corollary discharge, such as perisaccadic perceptual mislocalization, may be additionally modulated by the visual appearance of the saccade target. This is what we investigated here.

## Methods

### Experimental subjects and ethical approvals

We ran a perceptual mislocalization paradigm on 8 human subjects (4 female), aged 23-29 years. We also ran a control experiment on perceptual detectability of perisaccadic visual probes on 8 human subjects (5 female), aged 23-27 years. Four subjects were shared among the two paradigms, performing both variants of the experiments in different sessions.

The experiments were approved by ethical committees at the Medical Faculty of the University of Tübingen, and the subjects provided informed written consent. They were also financially compensated for their time, and the experiments were in line with the Declaration of Helsinki.

### Laboratory setup

The laboratory setup was the same as that used in our recent studies (14, 38), and we largely used similar procedures. Briefly, the subjects sat in a dark room 57 cm from a calibrated and linearized CRT display with 41 pixels/deg resolution. We tracked their eye movements at 1 KHz using a desk-mounted video-based eye tracker (EyeLink1000, SR Research), and we stabilized their head position using a custom-built apparatus described earlier (39). Stimuli were presented on a gray background of 20.01 cd/m^2^ luminance, and fixation spots (7 by 7 min arc) were white (with 49.25 cd/m^2^ luminance). The experiments were controlled using the Psychophysics Toolbox (40–42) with EyeLink Toolbox extensions (43).

### Experimental procedures

#### Perceptual mislocalization experiment

In the perisaccadic perceptual mislocalization experiment, the subjects fixated a white spot placed ∼8.5 deg either to the right or left of display center. After 500 ms, a low (0.5 cycles/deg; cpd) or high (5 cycles/deg; cpd) spatial frequency Gabor grating was presented at screen center. The grating had a gaussian smoothing window with σ=0.49 deg, and it had high contrast (100%); its phase was random. The subjects were still required to maintain fixation for another 1000-1500 ms (because we wanted to avoid any sensory transients associated with grating onset). When the fixation spot was removed, the subjects made a saccade towards the center of the grating, which had a central marker in its middle like in our neurophysiology experiments (37) (in order to minimize inter-trial saccade variance). Once we detected a saccade onset, as per the procedures we described recently (14, 38), we presented a brief (1-frame; 85 Hz refresh rate in the display) flash of a white square (that is, of high contrast) of size 45 by 45 min arc. The times of the flash were either immediately upon saccade detection or at +40 or +80 ms from online saccade detection. In post-hoc analyses, we recomputed the flash times to always report them relative to actual saccade onset (see Results). Moreover, the task of the subjects was to point (with a computer mouse) to the perceived location of the flash. Specifically, after the subjects fixated the central grating for 500 ms (that is, after the saccade), we removed all stimuli from the display, and we displayed a mouse cursor at display center. The subjects clicked on the perceived flash location, and they were free to move their eyes until they responded (they typically made a second saccade towards a region near the perceived flash position).

We picked four possible perisaccadic flash locations, each at a radius of ∼7.3 deg from the saccade target center location. One flash was directly opposite the saccade target direction (i.e. on the horizontal axis), and the other three flashes were along the saccade direction and ahead of the saccade target location: one on the horizontal axis directly ahead of the saccade target (along the same vector) and the other two at the +/-45 deg diagonals. Even though the flash opposite the saccade vector was expected to have the least mislocalization (17) (which we confirmed), we included it so that subjects could not expect always seeing flashes ahead of the saccade target. Nonetheless, in the analyses (see below), we only analyzed the three flash locations ahead of the saccade target. This is because the mislocalization for the backwards flash position was small in amplitude; past work that looked at mislocalization of the flash opposite the saccade vector typically used significantly larger saccades than we used, which made the smaller mislocalization of such a flash easier to detect than in our experiments (11, 16). In addition, and again because of our smaller saccade sizes than in previous studies, this backwards probe location was actually close to the initial fixation spot (only about 1 deg away from it when the eye was still fixated before saccade onset), which made us worry that there could be interactions with the initial fixation spot location in the subjects’ reported locations. Thus, it was more appropriate to analyze only the three forward probe flash locations.

Consistent with the above sentiment, we generally wanted to avoid having spatial references frames (e.g. the edges of the display monitor) strongly influence performance in the mislocalization task. Therefore, across trials, we randomly shifted all stimulus positions in a given trial by a random (but constant) number from the range of +/-1.2 deg in either horizontal or vertical direction (at pixel resolution; 1/41 deg). That is, on every trial, the overall displayed stimulus geometry was randomly shifted on the display monitor from previous trials, so that the monitor edges could not act (across trials) as reliable reference frames for probe flash localization. For example, if the initial fixation position was shifted by 1 deg to the right of its standard position on a given trial, then the grating position at the center of the display was also shifted by 1 deg to the right, and so was the perisaccadic probe flash (in other words, the stimulus relationships were all unchanged, but the overall absolute position was shifted by 1 deg to the right). This approach minimized the possibility of subjects remembering the absolute positions of flashes relative to the display monitor edges across trials. We are thus confident that the results that we report here were less affected by global biases of expected, world-centered, probe flash locations.

Finally, note that the low and high spatial frequencies chosen as the saccade targets were differentially represented by the SC motor bursts in our recent neurophysiological results (37). That is why we chose these particular spatial frequencies.

We analyzed 324 trials per spatial frequency in each subject.

#### Perceptual detection experiment

In a second control experiment, we tested the visibility of the perisaccadic probes used in the above experiment. That is, we wanted to confirm that the contrast used for the probe flashes in the experiment above was high enough to avoid effects of saccadic suppression. We also wanted to confirm that, even if suppression was avoided by the high probe contrasts, perceptual visibility for such high contrast probe flashes was still independent of the saccade target appearance. To do so, we collected full psychometric curves of probe detection (14). Thus, the experiment was the same as that described above, except that the probe flash now had variable contrast from trial to trial. Additionally, the probe flash time was either immediate upon online saccade detection, or at +11 or +35 ms. As we show in Results, this timing allowed us to investigate probe visibility in the two “compression” intervals in which we observed significant perisaccadic perceptual mislocalization in the main experiment above. Finally, in this task variant, the probe flash could appear at one of four cardinal locations relative to the saccade target center (and at an eccentricity halfway from the initial fixation position). The subjects’ task was to report which of the four locations had a brief flash at it, and they did so via a response button box (indicating that the flash was to the right of, left of, above, or below the saccade target location). Note that the eccentricity of the probe flashes used here was roughly similar to the perceived eccentricities that we report in Results for the mislocalization experiment, making the comparisons of the results in the two task variants meaningful.

Across trials, we varied the flash luminance to obtain full psychometric functions of flash detection sensitivity. The procedure was similar to that we described recently (14). Briefly, we performed a staircase procedure in the first session, to obtain an initial estimate of perceptual threshold in a subject; in subsequent sessions, we also added additional samples of flash contrast to sample more locations on the x-axis of the psychometric curves. Like in the mislocalization experiment above, we also randomized the starting position of the whole stimulus display items across trials. That is, a constant random shift was applied to all stimulus positions in any given trial.

We analyzed approximately 450-500 trials per spatial frequency in each subject.

### Data analysis

We detected saccades and microsaccades using our previously described methods (44, 45). Before analyzing any perceptual reports, we filtered the eye movement data to ensure matched saccade execution across low and high spatial frequency saccade targets. In the perceptual mislocalization task, we did so by binning the saccade landing points into bins of 0.5 x 0.5 deg. We also binned the saccade main sequence (46, 47) data points into bins of 0.5 deg x 40 deg/s. We then only accepted trials into the analyses that fulfilled the following criterion: they come from landing position and main sequence bins that had at least one trial repetition within them from each of the two saccade-target image appearances (low and high spatial frequencies). This way, the saccades across image appearances were matched for both metrics and kinematics (see Results). In the perceptual detection task, we used a slightly different, but equally effective (see Results), way of vector and kinematic matching. First, we defined a maximal radius (2 deg) for initial eye position at saccade triggering, and made sure that trials with both high and low spatial frequency saccade targets had the same initial starting position ranges. Similarly, we defined a maximal radius for final eye position at the end of the saccade, to make sure we had matched saccade vector ranges. Then, for each saccade direction (rightward or leftward), we ensured that the amplitudes and peak velocities of the saccades were matched regardless of whether the saccade target had a high or low spatial frequency (see Results). We also excluded trials with saccadic reaction times <80 ms or >500 ms.

To analyze perisaccadic perceptual mislocalization, we first confirmed that we had matched saccade vectors like described above. We also confirmed that saccade peak velocities (that is, kinematics) were similar for low and high spatial frequency saccade targets (see Results). We then recomputed all flash times relative to the actual saccade onset times after we properly detected the eye movements. After that, we collected each subject’s click locations at different flash times. We classified the flash times into three categories (t1, t2, and t3) according to the histograms of flash times that we observed after recomputing the saccade onsets from the recorded eye movement data (see Results). To summarize the subjects’ click locations, we first remapped all data from leftward saccades to reflect their results across the vertical axis. That is, we flipped the horizontal axis for leftward saccades, in order to pool all results with the rightward saccades. Thus, in Results, we always show observations on a schematic vector representation having rightward saccades. After such remapping, we picked the median click location for each subject, each flash probe time, each flash probe location, and each saccade-target appearance condition (low or high spatial frequency saccade target). Note that we always measured click locations relative to the stimulus locations; thus, the across-trial random shifts in global stimulus positions that we applied experimentally were removed in the analyses. To summarize the mislocalization effects, we measured for each trial the Euclidean distance from the actual flash location to the click location. In such summary analyses, we took (within a subject) the median Euclidean distances from all trials with forward flash positions; that is, for each subject we obtained a single (across-trial) Euclidean distance measure across the three probe flash locations that were ahead of the saccade target. Larger mislocalization for a given condition (e.g. one saccade image appearance) was associated with a larger Euclidean distance. We then plotted the mean and SEM of the distance across subjects, and we showed descriptive statistics and underlying individual subject results. We report all the statistical test in Results.

We also looked at the direction of the mislocalization vector for oblique probe flash positions. That is, mislocalization is known to have a two-dimensional landscape (along saccade direction and orthogonal to it) even if the saccades themselves are cardinal (16, 17). This two-dimensional landscape can be investigated by measuring the component of mislocalization independently either parallel to the saccade direction vector or orthogonal to it. Thus, we picked the two oblique flash positions that were ahead of the saccade target (at the +/-45 deg diagonals), and that were expected to have the strongest oblique mislocalization (16, 17). Then, we compared how either their parallel or orthogonal mislocalization was different between the two image appearances of the saccade targets. For the parallel component, we measured the horizontal component of their Euclidean distance measure described above, and for the orthogonal component, we measured the vertical component.

Finally, we compared mislocalization for flashes presented in the upper versus lower visual fields. That is, it is well established that the SC has an asymmetric representation of the upper and lower visual fields, which has been shown to play a potential role in the influence of saccade direction and visual field location on the strength of both perceptual mislocalization (17) and perceptual saccadic suppression (48). We, therefore, asked whether such an asymmetry could also differentially affect mislocalization strengths for upper and lower visual field flashes even with the same saccade direction. We did this by taking only one image condition (0.5 cycles/deg) and analyzing mislocalization strength for either the oblique flash that was presented in the upper visual field or the one that was presented in the lower visual field. Very similar results were obtained with the high spatial frequency saccade target, so we only reported the low spatial frequency result for simplicity.

All statistics on the percpetual mislocalization data were performed with MATLAB (Version R2020b) and IBM SPSS Statistics (Version 29.0).

In the perceptual detection experiment, we obtained psychometric curves of perceptual performance as a function of probe contrast level. To do that, we used the Psignifit 4 toolbox (49) for each subject individually, using a cumulative gaussian function (and 0 lapse rate). We then averaged across the subjects’ individual psychometric curves to obtain a population measure; thus, the population psychometric curve was the average of 8 individual psychometric curves. We did this for each time bin of flash times relative to saccade onset (with the time bins defined as t1a, t1b, and t2, respectively; see Results). Statistically, we performed bootstrapping to check whether perceptual detection depended on the saccade target image type. To do so, we created random permutations (10000) in which we randomly picked trials for the two conditions (low or high spatial frequency saccade target) within each subject. When doing so, we kept the same numbers of observations per subject per condition, in order to maintain the sampling numbers consistent with our actual experimental data (especially after eye movement filtering). We then performed psychometric curve fits from such permuted data and measured the threshold at the 62.5% correct rate. Across the 10000 permutations, we plotted the distributions of thresholds between the two sets of permuted conditions. If our actual measured threshold difference between low and high spatial frequency saccade targets was deviated by more than 95% of the threshold differences of the random permutations, then we deemed our measured threshold difference in the real experiment to be significant.

## Results

Our aim was to explore whether the strength of perisaccadic perceptual mislocalization could be modulated by the visual appearance of saccade targets, as might be predicted from observations of sensory tuning in SC saccade-related motor bursts (37). To do so, we designed an experiment in which humans reported the location of a brief perisaccadic probe flash that was presented at different times relative to saccade onset. Specifically, each trial consisted of an instruction to generate a saccade towards the center of a Gabor grating located near the center of the display screen (Fig. 1A). The initial fixation position was 8.5 deg to either the right or left of the grating center, thus requiring horizontal saccades. In addition, the Gabor grating could have one of two different spatial frequencies (Methods). At different times relative to the online detection of a saccade (Methods), we presented a brief, single-frame probe flash at one of four locations relative to the saccade target location. Three of these possible flashes were ahead of the saccade target, and one was behind it (Fig. 1A; Methods).

Figure 1B shows the distribution of probe flash times that we sampled for one example subject. During actual data collection, the computer triggered the probe flash at one of three possible delays after online saccade detection. However, the real delay was necessarily different since online saccade detection required some data buffering for eye speed estimations, and since there was some variance in the online speed estimates. Therefore, in post-hoc analyses, we re-detected all saccades (Methods), and we then plotted the distributions of flash times in Fig. 1B. As can be seen, relative to the generated saccades (Fig. 1C, D, bottom shows radial eye speed from the saccades of the same subject), one set of times was intra-saccadic (labeled t1), another was shortly after the saccades (labeled t2), and the third was even longer after the saccades, when perception was expected to be nearer to a veridical performance state (labeled t3). Thus, our paradigm allowed us to analyze periods of significant perisaccadic mislocalization (times t1 and t2) as well as periods closer to full perceptual recovery (t3). For simplicity, in what follows, we sometimes refer to t1, t2, and t3 below as 30 ms, 70 ms, and 110 ms, respectively.

Importantly, we ensured that the saccade vectors and kinematics were matched for the two different saccade target appearances. This was aided by having a small target spot in the middle of the gratings (37) (Fig. 1A), as well as by our filtering criteria described in Methods. The net result was that the saccade distributions for the two different image types were virtually indistinguishable from each other between the low and high spatial frequency saccade targets (Fig. 1C, D; one example subject’s results are shown). Thus, we were now in a position to explore whether perceptual mislocalization was different for the two different saccade target images, despite the matched saccades.

**Figure 1.**
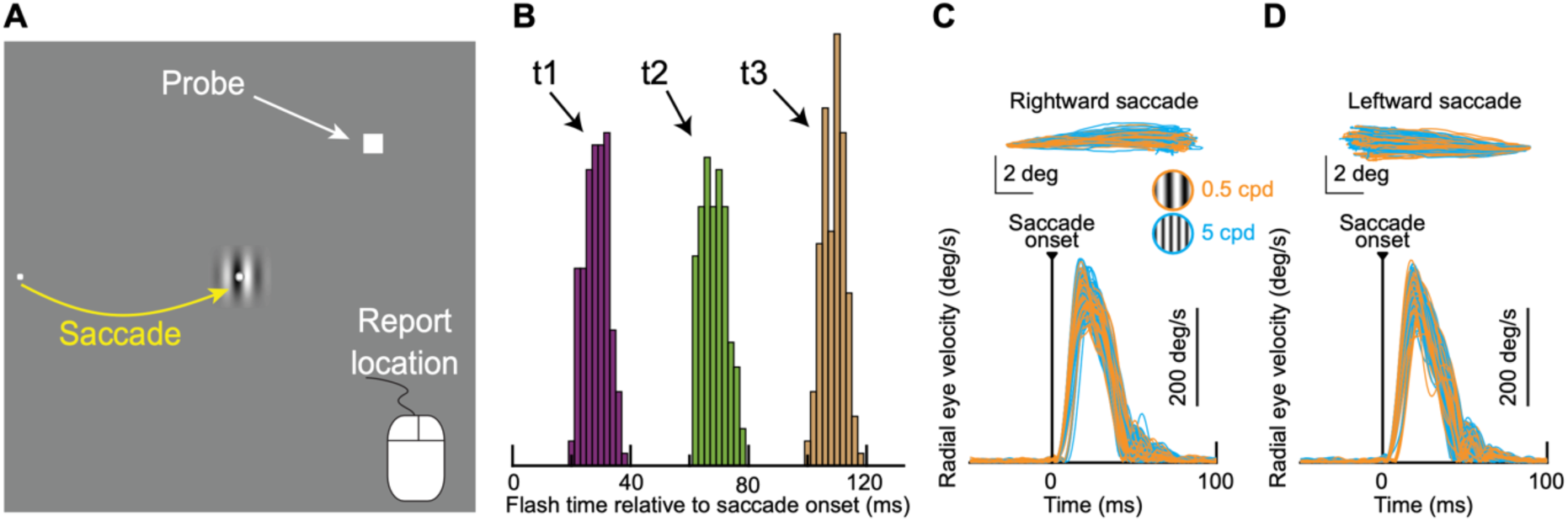
Probing perisaccadic perceptual mislocalization with different saccade-target image appearances. **(A)** Subjects generated a rightward (shown) or leftward saccade towards the center of a low or high spatial frequency grating. The center of the grating always had a small spot, in order to improve saccade accuracy/precision and vector matching across different trials and different saccade-target appearances (Methods). At different times relative to online saccade detection, we presented a very brief probe flash at one of four positions. Later, all stimuli were removed, and subjects had to point to the perceived flash position using a mouse cursor. **(B)** Distribution of probe flash times relative to saccade onset in one example subject. We classified flash times according to the three seen peaks, and we labeled the categories t1, t2, and t3, respectively. We expected perceptual mislocalization to progressively get smaller and smaller from t1 to t3 (11). N=460 trials. **(C)** The top panel shows horizontal and vertical eye position traces for rightward saccades from an example subject. Starting eye positions from all trials were aligned for better visualization of the saccade vectors, and the different colors show saccades towards either the low or high spatial frequency grating. The bottom panel shows radial eye speed traces from the same trials. Note how saccade metrics (top) and kinematics (bottom) were matched across image types. Also note how the flash times in **B** spanned intra-saccadic flashes (t1) as well as flashes immediately after (t2) or later well beyond saccade end (t3). N=113 and 117 for low and high spatial frequency saccade targets, respectively. **(D)** Same as **C** but for leftward saccades from the same subject. N=117 and 113 for low and high spatial frequency saccade targets, respectively.

### Stronger perisaccadic mislocalization for low spatial frequency saccade targets

We analyzed each subject’s click positions in each condition and time bin from Fig. 1. As stated in Methods, we focused on the three probe flash locations ahead of the saccade target (one directly along the saccade vector and two at +/-45 deg oblique directions). This was the case especially because these three locations cause the strongest mislocalization (16, 17). Moreover, given that our saccades were significantly smaller than in previous studies of mislocalization (11, 16) (and were thus associated with smaller mislocalization strengths in general), and given the proximity of the probe opposite the saccade vector to the initial fixation position (Methods), we felt that focusing only on the probe flash locations ahead of the saccade target was justified.

In Fig. 2A-C, we show all click positions from one example subject’s data. Each panel shows results from a given time bin, and this subject is the same subject whose eye movement data are shown in Fig. 1. As can be seen, the subject exhibited clear perisaccadic perceptual mislocalization, which recovered with increasing time from saccade onset. That is, regardless of which saccade target was visible (the two different colors in Fig. 2), the subject’s click positions were closer to the saccade target in Fig. 2A than they were in Fig. 2B, C. And, by time t3 (110 ms), the click positions were closer to being veridical and near the true probe flash positions (Fig. 2C) (with the existence of some expected residual systematic undershoot biases; refs. (50, 51)).

Importantly, Fig. 2A-C shows that the subject’s perceptual reports clearly depended on the appearance of the saccade target. This is seen through the different colors of the clicks in Fig. 2A-C. In orange, we show the clicks with the low spatial frequency saccade target, and in blue, we show clicks with the high spatial frequency saccade target. There was stronger mislocalization, especially for the oblique probe flash positions, with the low spatial frequency saccade target. This was also evident in the median click positions for the different probe flash locations and times (Fig. 2D-F; error bars denote inter-quartile ranges). Thus, in this example subject, perisaccadic perceptual mislocalization depended on the visual appearance of the saccade target, even with metrically and kinematically matched saccades.

**Figure 2.**
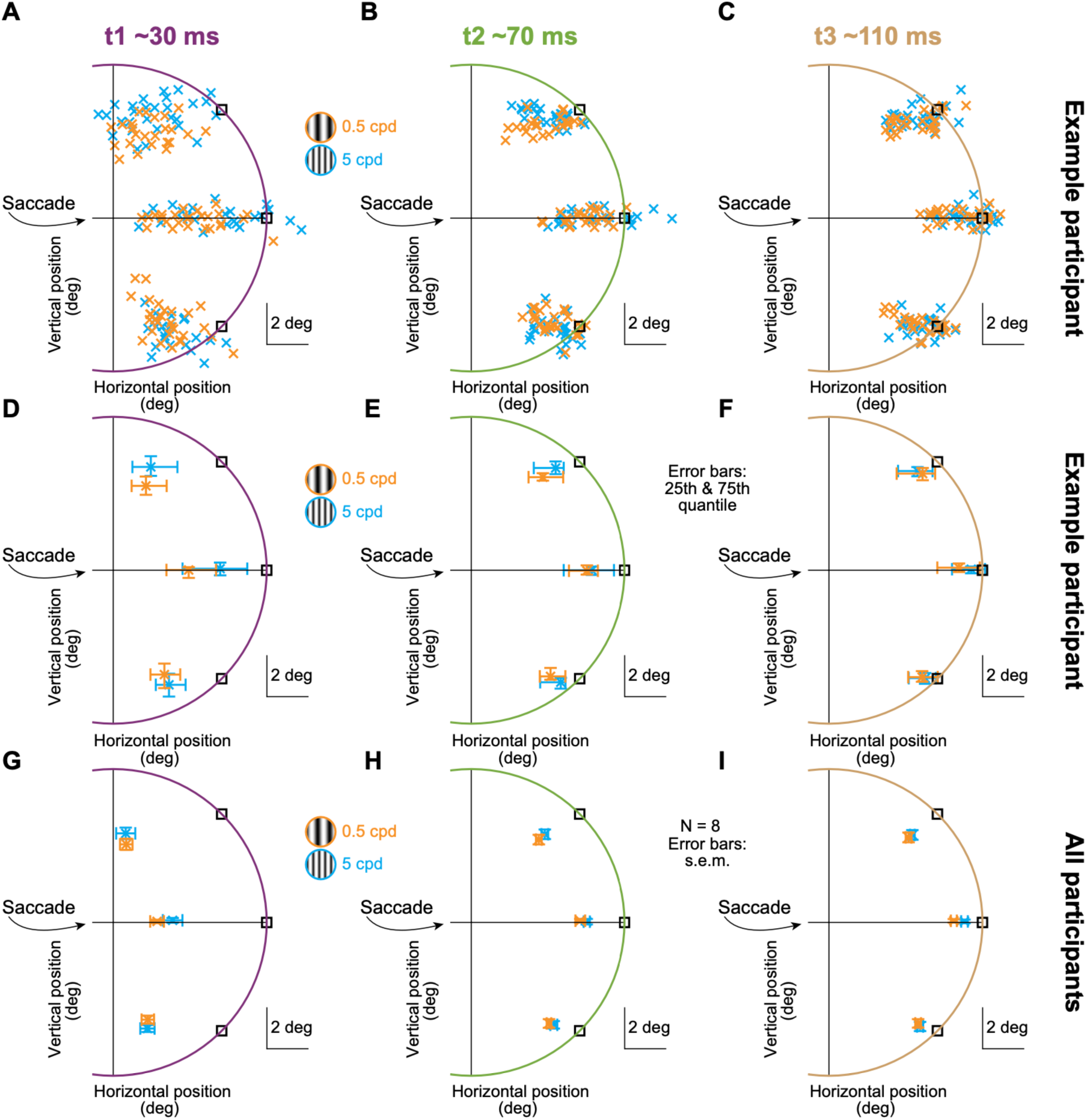
Stronger perisaccadic perceptual mislocalization with low spatial frequency saccade targets. **(A-C)** All click positions from one example subject (the same as in Fig. 1) after remapping all data to a rightward saccade direction (Methods). The different panels show results from when the probe flashes were presented at time t1 (**A**), t2 (**B**), and t3 (**C**), respectively, relative to saccade onset. Mislocalization got progressively weaker with time (compare the panels). Importantly, the mislocalization within a given time bin was different for the two different saccade-target image appearances (compare the different colors in each panel). N=80, 73, 77 trials for the clicks with the low spatial frequency saccade target in **A**, **B**, and **C**, respectively. N=75, 77, 78 trials for the clicks with the high spatial frequency saccade target in **A**, **B**, and **C**, respectively. **(D-F)** Each symbol shows the median click location of the same subject, for a given probe flash location, time, and saccade-target image appearance. The range bars around each median click location indicate the inter-quartile extents of the data. **(G-I)** For all analyses like in **D**-**F**, we averaged across 8 subjects. Error bars denote SEM across the subjects. Also see Figs. 3, 4 for more detailed quantitative analyses.

These results were consistent across our 8 subjects. We repeated the analyses of Fig. 2D-F for each subject, and we then plotted the mean and SEM values across the subjects in Fig. 2G-I. There were systematic differences between the two different saccade-target image appearances, which we investigated further in Fig. 3. Specifically, we computed, for each trial, the Euclidean distance between the true probe flash position and the individual click position. Then, within each subject, we took the median distance across all three flash locations ahead of the saccade target location, and we did this separately for the different probe flash times. The saturated data points in Fig. 3 show the mean and SEM across 8 subjects, and the gray data points show individual subject results. Note that the y-axis ranges in Fig. 3A-C are not the same across the three panels, and this is because mislocalization got progressively smaller with increasing time (Fig. 2), and we wanted to zoom into the data ranges of each panel individually. As can be seen, for each of the times t1 and t2, all but one subject showed a stronger perisaccadic perceptual mislocalization for the low spatial frequency saccade target than for the high spatial frequency saccade target (Fig. 3A, B). At time t3, all but two subjects still showed this effect, although the overall mislocalization strength was much weaker given the longer time after the saccade (Fig. 2).

We further quantified these effects by plotting in Fig. 3D the difference in the mislocalization strength between the low and high spatial frequency saccade target images as a function of probe flash time. As can be seen, this difference was similar for times t1 and t2, and it was clearly positive in each of these two time points (with only one outlier subject in each case). Statistically, we performed a repeated measures two-way ANOVA with the factors time and spatial frequency of the saccade target. Both main effects were significant (time: F(2, 14)=67.78, p<10^-7^, η^2^=0.906; spatial frequency of the saccade target: F(1, 7)=69.04, p<10^-4^, η^2^=0.908). There was no significant interaction between these two factors. This is consistent with the observation that the difference in mislocalization strength for the two different saccade target image appearances was very similar for times t1 and t2 in Fig. 3D. We also performed post-hoc tests (with Bonferroni correction), which revealed a significant difference in mislocalization strength between the saccade target image types for both time t1 (p=0.015) and time t2 (p=0.025). These results suggest that a perisaccadic perceptual phenomenon, which can potentially depend on ascending SC projections (27–30), depends on the visual appearance of the saccade target (especially at times t1 and t2). This implies that sensory tuning in SC neuronal movement commands (37) may, in turn, potentially provide more than just the vector information of executed movements via SC-sourced corollary discharge signals.

**Figure 3.**
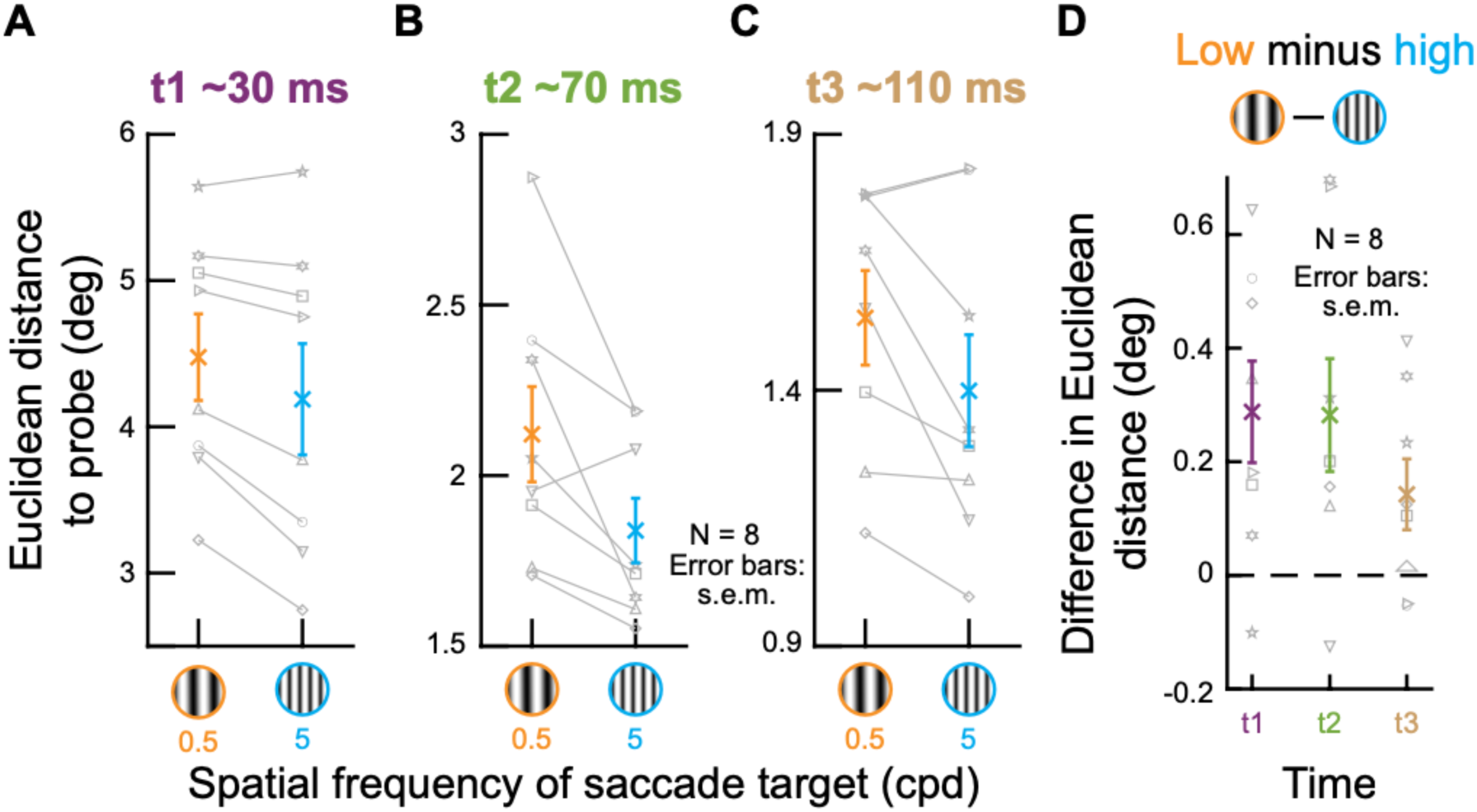
Population summaries of the results in Fig. 2. **(A)** The saturated data points show the mean and SEM across subjects (N=8) of the Euclidean distances between the median click location at time t1 and the true probe flash location (for the three probes ahead of the saccade target; Fig. 2). The gray data points show individual subject results. All but one subject had stronger perisaccadic perceptual mislocalization for the low spatial frequency saccade target. **(B)** Similar results for time t2. Again, all but one subject had stronger perisaccadic perceptual mislocalization for the low spatial frequency saccade target. **(C)** Similar results for t3. Here, all but two subjects showed stronger mislocalization for the low spatial frequency saccade target. Note that the y-axes are different in the different panels due to the time varying amplitude of the mislocalization effect (Fig. 2). **(D)** At each flash time (x-axis values), we plotted the difference in mislocalization strength between the low and high spatial frequency saccade target image trials. A positive difference indicates stronger mislocalization for the low spatial frequency saccade target. The saturated data points show the mean and SEM across subjects, and the gray data points show the individual subject results. Especially at times t1 and t2 (with strong perceptual mislocalization), there was stronger mislocalization for the low spatial frequency saccade target appearance. Also see Fig. 4.

### Stronger orthogonal mislocalization for low spatial frequency saccade targets

In the analyses of Fig. 2, we additionally noticed that oblique probe flash positions exhibited a stronger dependence of mislocalization strength on saccade-target image appearance in the direction orthogonal to the saccade vector than in the direction parallel to it. Specifically, at time t1, with maximal mislocalization strengths in our experiments, both the example subject (Fig. 2A, D) and the population (Fig. 2G) showed a bigger deviation in the vector direction between the true probe flash position and the perceptual reports of the subjects, suggesting a difference in the orthogonal component of mislocalization. Such a difference was masked by the analyses of Fig. 3, which pooled different probe flash positions, and which were somewhat agnostic of the two-dimensional nature of perceptual mislocalization. Therefore, to explore this further, we expressed the mislocalization effect according to its two orthogonal spatial components: one along the direction of the saccade (parallel to it) and one orthogonal to the saccade vector (Fig. 4A). We then compared the mislocalization effect in each direction separately for the two different saccade-target image appearances.

**Figure 4.**
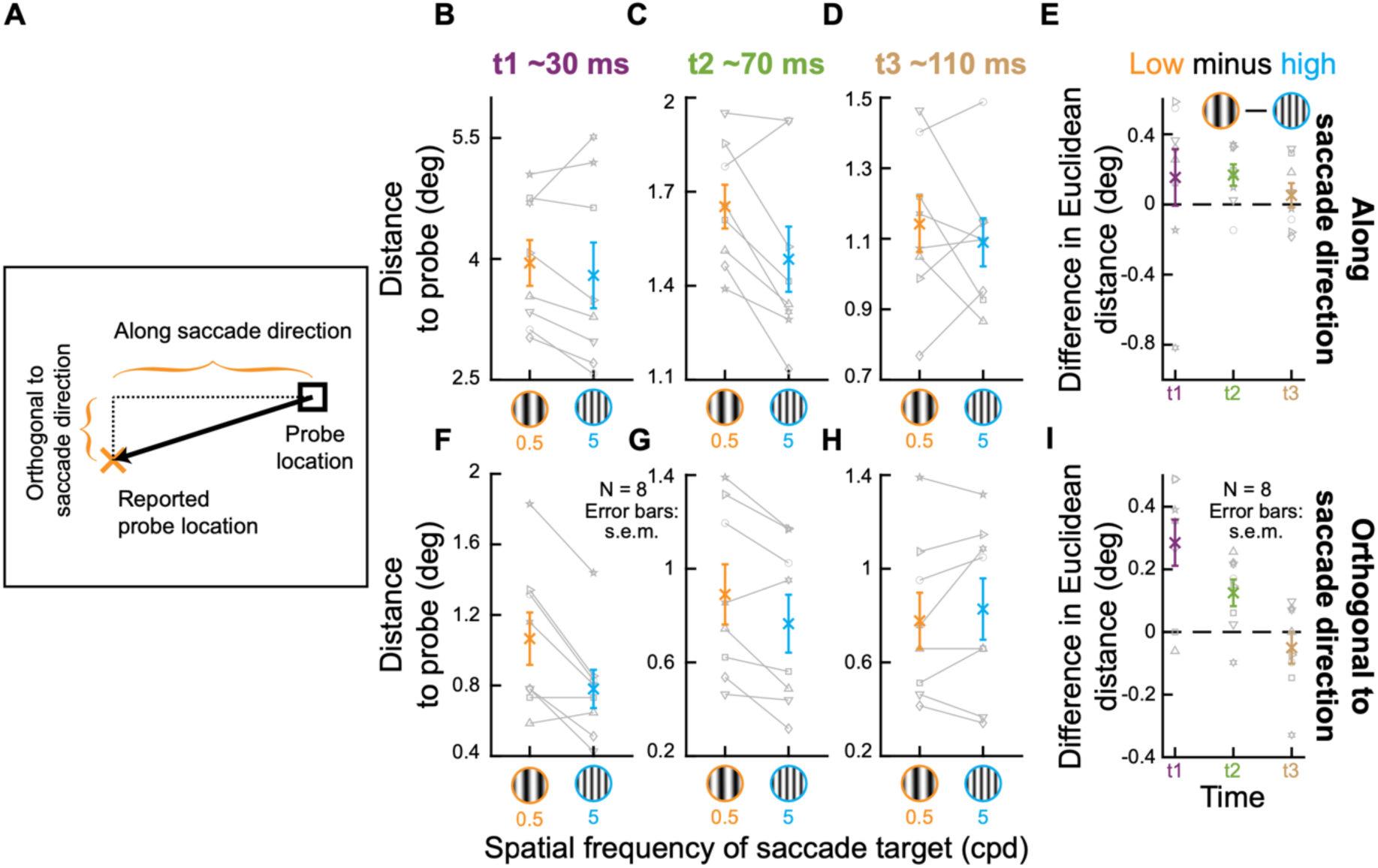
Stronger orthogonal mislocalization with low spatial frequency saccade targets. **(A)** Our analysis approach for assessing the strength of orthogonal mislocalization. For the oblique probe flash positions ahead of the saccade target, we measured the two components of mislocalization separately. One component was along the saccade direction (measuring the horizontal distance between the reported and true probe locations), and one component was orthogonal to the saccade direction (measuring the distance between true and reported probe locations along the orthogonal axis). **(B-E)** Same as Fig. 3A-D but for the parallel component of mislocalization. The trends were all the same as in Fig. 3, but the significant effect of saccade-target image appearance only emerged at time t2 (see text). This might suggest that orthogonal mislocalization might have dominated the effects of Fig. 3, especially at time t1. **(F-I)** We confirmed this by repeating the same analyses, but this time for the orthogonal component of mislocalization. Note how there was clearly stronger orthogonal mislocalization for low than high spatial frequency saccade targets, especially at time t1. All other conventions in this figure are the same as those in Fig. 3.

We first compared the mislocalization effects along the saccade direction (Fig. 4B-E), which is generally the direction in which mislocalization is strongest even for oblique flash positions (16, 17). While the trends were all the same as those seen in Fig. 3 above, the clearest statistical effect now emerged only at time t2. Specifically, we performed a repeated measures two-way ANOVA with the factors time and spatial frequency of the saccade target. Both main effects were significant (time: F(2, 14)=47.27, p<10^-6^, η^2^=0.871; spatial frequency of the saccade target: F(1, 7)=6.6, p=0.037, η^2^=0.486), and there was no interaction. However, in post-hoc tests (with Bonferroni correction), there was only a significant difference for the two types of saccade target at time t2 (Fig 4C, E). Thus, this might indicate that the saccade-target image appearance effects seen in Fig. 3, especially at maximal mislocalization periods (time t1), might have been dominated by the orthogonal component of mislocalization. We confirmed this to be the case. We performed the same tests but now on mislocalization orthogonal to the saccade direction (Fig 4F-I). Again both main effects were significant (time: F(2, 14)=4.34, p=0.034, η^2^=0.383; spatial frequency of the saccade target: F(1, 7)=11.12, p=0.013, η^2^=0.614). On top of that, there was also an interaction between the factors time and spatial frequency of the saccade target (F(2,14)=9.47, p=0.003, η^2^=0.575), indicating that the difference in mislocalization for the different saccade targets differed as a function of time. Consistent with this, a post-hoc test (with Bonferroni correction) revealed that the significant difference as a function of saccade-target image appearance was present at both times t1 (p=0.006) as well as t2 (p=0.022) (Fig 4I). Thus, even the two-dimensional landscape of perisaccadic perceptual mislocalization (16, 17) depends on the saccade-target image appearance, and this effect was clearest at time t1 (Fig. 4I).

### Stronger mislocalization for probes in the upper rather than lower visual field

The SC is known for its asymmetric representation of the upper and lower visual fields (21, 22). Therefore, we also wondered whether, even for purely horizontal saccades, mislocalization strength could additionally depend on the visual field locations of the probe flashes. For example, it could be the case that upper visual field magnification in the SC (21) is an additional topographic map distortion that can potentially modify mislocalization patterns. At least according to accounts of perceptual mislocalization that depend on anatomically distorted topographic maps (19), this is a distinct possibility. Testing this possibility would provide even further motivation for the idea of a putative involvement of the SC in remapping mechanisms that can cause perisaccadic perceptual mislocalization. Moreover, if validated, showing different perisaccadic mislocalization for the upper and lower visual fields would add to increasing evidence that perisaccadic vision in general might follow SC visual field asymmetries rather than asymmetries in perceptual performance that occur in the absence of saccades (likely cortically mediated), which are of exactly opposite sign to the SC asymmetries (52–55). Indeed, this is already known to be the case for perisaccadic suppression of visual sensitivity (48).

For the low spatial frequency grating as the saccade target, we compared mislocalization strength when the probe was presented in either the upper or lower visual field. That is, we explored the two oblique probe locations ahead of the saccade target location, which were perfectly symmetric with respect to the saccade vector, except for the visual field difference (Methods). The results are shown in Fig. 5, which is formatted similarly to Figs. 3, 4 above. However, here, instead of comparing the effects of the saccade-target image appearance on mislocalization, we now compared probe flash locations for a single saccade target type. As can be seen, there was clearly stronger mislocalization in the upper, rather than lower, visual field. We tested this statistically with a repeated measures two-way ANOVA (factors: time and probe location). The test results were significant for both the factors of time (F(2, 14)=73.66, p<10^-7^, η^2^=0.913) and probe location (F(1, 7)=30.61, p<10^-3^, η^2^=0.814). There was also a significant interaction between these two factors (F(2,14)=5.52, p=0.017, η^2^=0.441) (Fig. 5D). Moreover, post-hoc tests revealed that the difference between upper and lower visual field probe locations was significant at all three tested times (t1: p<0.001; t2: p=0.002; t3: p=0.021) (Fig 5D). Therefore, there was a clear dependence of perisaccadic perceptual mislocalization strength on visual field location, even with purely horizontal saccades. Note that we reached similar conclusions when we performed these analyses for the high spatial frequency saccade targets as well (this can also be easily inferred from Fig. 2).

**Figure 5.**
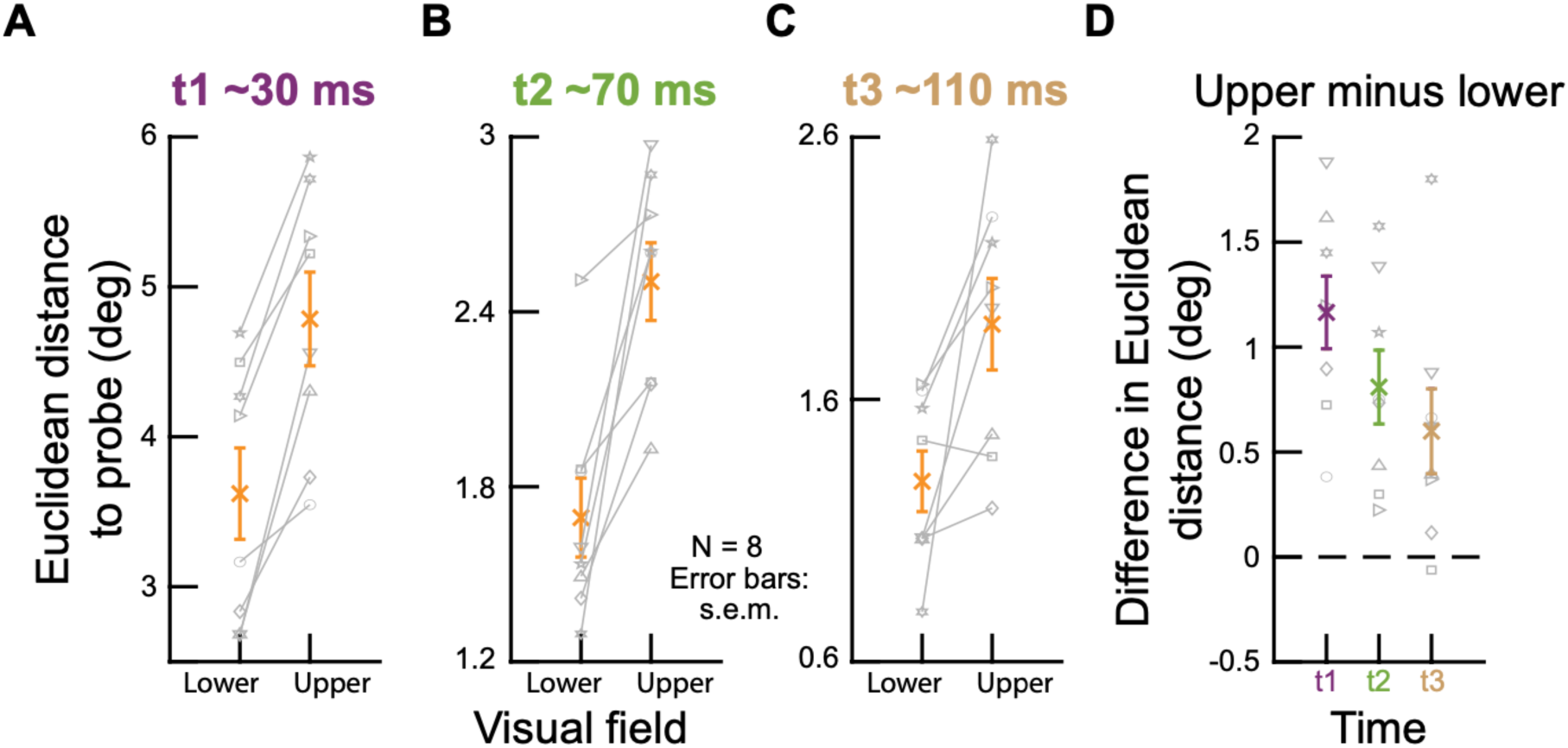
Stronger perisaccadic perceptual mislocalization for probe flashes in the upper, rather than lower, visual field. This figure is formatted identically to Fig. 3, except that the comparison is now between visual field locations rather than saccade-target image appearances. Specifically, we only considered the trials with 0.5 cycles/deg spatial frequency saccade targets. Then, we considered the mislocalization strength of the two probe locations that were in oblique directions relative to the saccade target. Because of the configuration of our display, these two probe locations were at identical positions relative to the saccade target (Method). However, one of them was in the upper visual field, and the other was in the lower visual field. As can be seen, there was stronger mislocalization for upper visual field probes. All other conventions are identical to those in Fig. 3.

Thus, our results so far indicate that perisaccadic perceptual mislocalization depends on both saccade-target image appearance (Figs. 2-4) and retinotopic probe flash location (Fig. 5). We next confirmed that these observations could not be explained by an altered visibility of the probe flashes by the visual conditions of our experiments.

### Similar flash visibility for low and high spatial frequency saccade targets

In the above experiment and analyses, we were confident that the probe flashes were supra-threshold (and thus highly visible to the subjects) because of their high contrast (Methods). Therefore, we interpreted the results as suggesting that perceptual mislocalization can indeed depend on the visual appearance of the saccade target (Figs. 1-4). However, it could still be the case that the appearance of the saccade target (especially given that it was a relatively big image patch) could, in one way or another, cause different detectability levels of the brief perisaccadic probe flashes. Therefore, in a second experiment, we explicitly characterized the perceptual detectability of the flashes at different contrast levels. This allowed us to confirm that the detectability of the high contrast probe flashes in the results above was the same for either the low or high spatial frequency saccade targets.

We perisaccadically flashed brief, low contrast probes (Methods), and we measured perceptual contrast sensitivity curves. The paradigm itself was very similar to that used above. However, instead of localizing probe flashes, the subjects knew in advance that the probe could appear at one of the four cardinal directions around the saccade target (Fig. 6A; Methods). They simply had to report which of the four locations displayed the flash.

Like in the perceptual mislocalization experiment, we also had variable probe flash times relative to saccade onset. Specifically, Fig. 6B shows the timing of the probe flashes in this new experiment. As can be seen, the trigger points of the probe flashes after online saccade detection (Methods) resulted in a bimodal distribution of flash times. The second mode (at around 70 ms; Fig. 6B) was very similar to that in t2 of the original experiment. Since time t2 in that experiment was still a time in which we had clear differential mislocalization performance for the two different saccade-target image appearances (Figs. 2-5), this time bin in the current experiment was ideal to check whether the perceptual detectability of the probe flashes was any different for the two different saccade-target image features. We thus labeled this second mode in Fig. 6B as t2, since it was quantitatively similar to time t2 in the original experiment. As for the first mode in the histogram of Fig. 6A, we split it into two sub-categories (t1a and t1b) because we wanted to check for a worst-case scenario about potential visibility differences with maximal saccadic suppression, which would be expected for t1a (the closest time to saccade onset).

**Figure 6.**
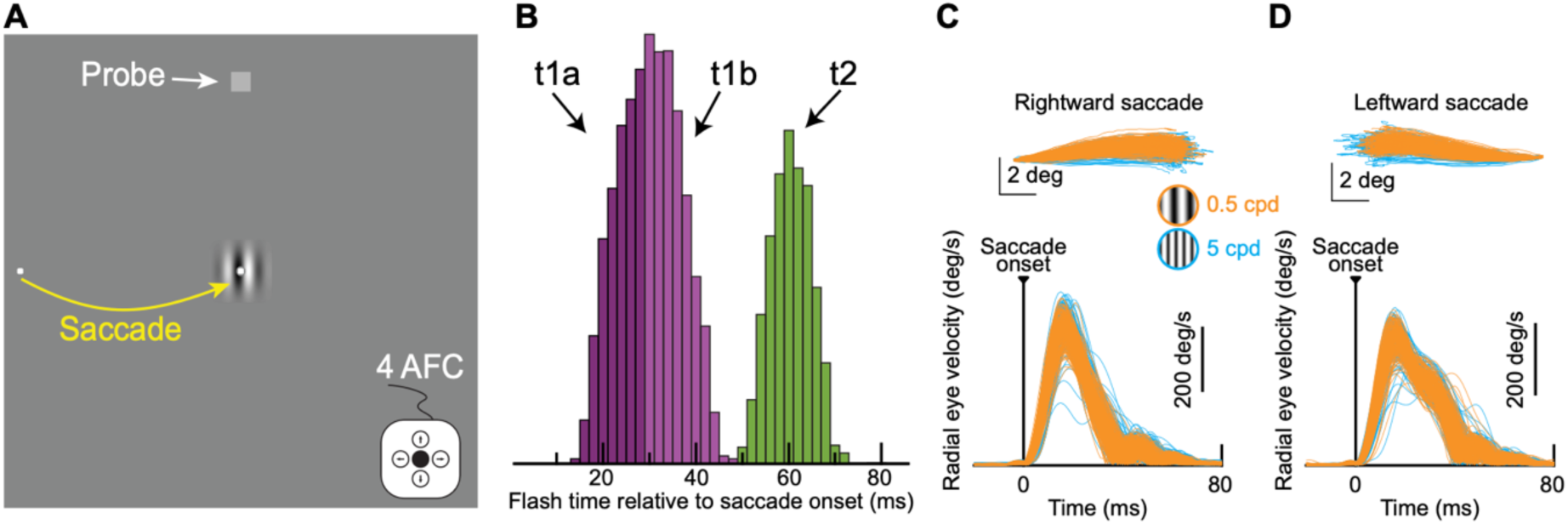
Probing perisaccadic perceptual detectability with different saccade-target image appearances. **(A)** Subjects generated a rightward (shown) or leftward saccade towards the center of a low or high spatial frequency grating. At different times relative to online saccade detection, we presented a very brief probe flash at one of four cardinal positions (pure up shown). Subjects had to indicate at which one of the four locations they detected the stimulus (4-alternative forced choice, 4 AFC, task). **(B)** Distribution of flash times relative to saccade onset in our data, from a sample subject. We classified flash times according to the three seen categorizations, and we labeled the categories t1a, t1b, and t2, respectively. t2 was quantitatively similar to t2 in the perceptual mislocalization experiment above. **(C, D)** Similar analyses to Fig. 1C, D, demonstrating that we matched saccades both in terms of their metrics and kinematics across the two saccade-target image appearances.

In the current experiment, we also matched the saccade metrics and kinematics across the two different image appearances of the saccade target, just like we did in the perisaccadic perceptual mislocalization experiment above. For example, Fig. 6C, D shows eye movement analyses very similar to those in Fig. 1C, D from a sample subject. As can be seen, the saccade properties were well matched across the two different image appearances of the saccade target. Therefore, we could now check the psychometric curves.

Even under the worst-case scenario of maximal saccadic suppression, perceptual detection reached ceiling performance at probe luminances well below those that we used in the perisaccadic perceptual mislocalization experiment described above. This can be seen from Fig. 7. In Fig. 7A, we show the psychometric curves of performance from one example subject (the same one as in Fig. 6C, D). One pair of curves is for probe flashes occurring at time t1a from Fig. 6B, and another pair is for probe flashes occurring at time t2. In each pair, one curve is for the low spatial frequency saccade target, and the other is for the high spatial frequency saccade target. For reference, the probe contrast used in the perisaccadic mislocalization experiment above was at an x-axis value of 70 in these psychometric curves. Thus, in all cases, this subject’s perceptual performance reached ceiling performance for much lower probe contrasts, even at the worst-case scenario of near-maximal saccadic suppression. Importantly, at time t2, when mislocalization still showed significant differential effects between saccade-target appearances (Figs. 2-4), the full psychometric curves in this current experiment were completely overlapping (and with very low detection thresholds). Thus, the results of Figs. 2-4 above cannot be explained by different probe detectability due to the different saccade-target image appearances used in the experiments.

These results were consistent across all 8 subjects (Fig. 7B). Interestingly, at the population level, we found a small, but significant (p<0.05; bootstrapping; Methods), difference in the semisaturation contrasts of the two saccade-target image appearances only at time t1a (but not at times t1b and t2). This might suggest that even perisaccadic perceptual suppression itself, which can putatively also rely on corollary discharge information (27, 56), might also depend on the saccade-target visual features. This is consistent with both the predictions of the neurophysiology (37) as well as with the general motivations for the current study. Nonetheless, from the perspective of perisaccadic perceptual mislocalization, which is the main topic of the current study, the most important feature of the results of Fig. 7B is that, at the probe contrasts used in Figs. 1-5 (x-axis luminance of 70 in Fig. 7B), perceptual detectability of the probe flashes was clearly at ceiling and did not depend on the saccade-target image appearance. Thus, the results of Figs. 2-4 cannot be explained by visual-visual interactions caused by the different image appearances of the low and high spatial frequency saccade-target gratings.

**Figure 7.**
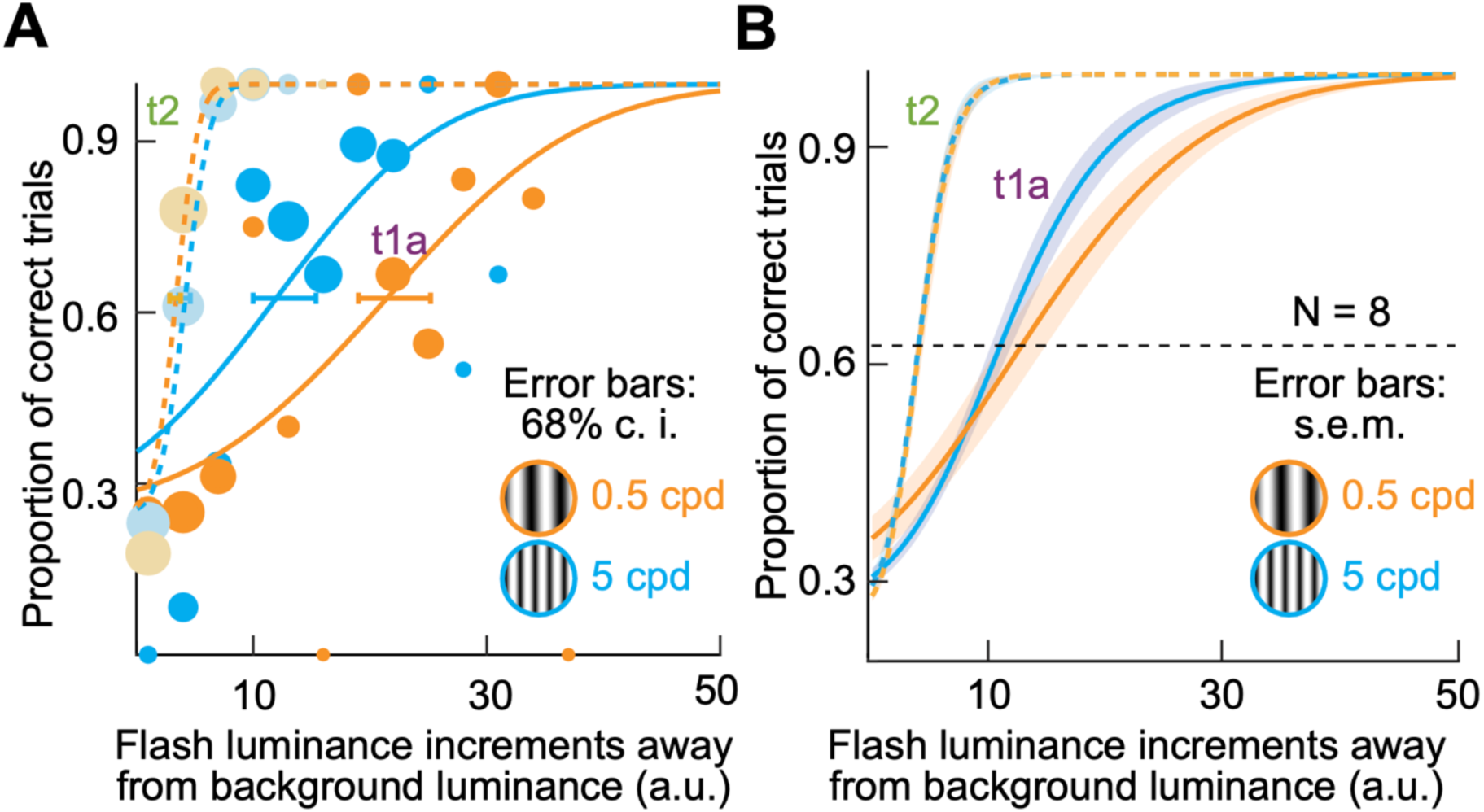
Dependence of probe detection on the saccade-target image appearance, but only for very low contrast probes and only during maximal saccadic suppression. **(A)** Psychometric curves of perceptual detection performance from one example subject. Since this was a four-alternative forced choice task, chance performance was at a 25% correctness rate, and threshold was defined as the contrast giving rise to 62.5% correctness rate. When the probe appeared during time t1a in Fig. 6B, the subject showed a higher detection threshold for the low spatial frequency saccade target than for the high spatial frequency saccade target. However, note that ceiling performance was reached by luminance index 50 above background. For reference, the probe luminance in Figs. 1-5 was at 70 on the x-axis. Thus, even at time t1a, the probe in Figs. 1-5 was highly detectable. For time t2, the thresholds were much lower, and the psychometric curves were almost completely overlapping. Note that at this time, perceptual mislocalization still significantly differentiated between the saccade-target image appearances (Figs. 2-4), suggesting that such differentiation was not mediated by different probe visibility for the two different types of saccade targets that we used. Error bars denote 68% confidence intervals. The size of each shown data point is scaled by the numbers of observations. **(B)** Population results. Each curve shows the average of the 8 subjects’ psychometric curves, and error bars denote SEM. At time t1a, there was a small threshold difference between the different saccade-target appearances (p<0.05; bootstrapping; Methods). However, at t2 (and t1b; not shown), there were no differences. Importantly, performance was clearly at ceiling at the probe contrasts used in the mislocalization experiment. Thus, the results of Figs. 2-5 cannot be explained by altered detectability of the probe flashes by the different saccade-target image appearances.

Therefore, our results, combined, suggest that perisaccadic perceptual mislocalization seems to depend on the visual appearance of the saccade target (Figs. 1-4), and that this dependence is not explained by a simple visual interaction between such appearance and the ability to detect high-contrast perisaccadic probe flashes (Figs. 6, 7).

## Discussion

In this study, we were motivated by our recent observations that SC motor bursts can be different for different visual appearances of the saccade target, even with matched saccade metrics and kinematics (37). Such difference in SC saccade-related discharge suggests that the motor bursts in the SC might be dissociated from the exact moment-to-moment execution of the eye movements (22). If so, then what function could such sensory tuning in SC motor bursts have? We hypothesize that it can be functionally used via the ascending projections from the SC to other brain areas. Such projections may be thought of as corollary discharge (23, 30): neurons on the SC map that are active at the time of saccade onset indicate the vector of the intended saccade (57), and such vector information (and the concomitant timing of the bursts) could help to either suppress visual sensitivity (28, 58) or remap retinotopic response fields (30) in the cortex. However, the SC benefits from integrating a large amount of visual information from many brain areas, including from the retina (59). Thus, at the time of saccade generation, visual information in the motor bursts could additionally allow the predictive processes associated with corollary discharge (28, 30, 60) to be more general than just informing cortex of the timing and vector of saccades. These processes could additionally allow trans-saccadic feature prediction. In other words, the SC could relay its own evidence of the visual appearance of the peripheral saccade target, and this evidence could potentially be useful for postsaccadic visual processing in the fovea. While this idea needs to be explicitly tested neurophysiologically (for example, by investigating the transfer of visual information to foveal neurons in the SC and elsewhere across saccades), our goal here was to check whether perceptual correlates of it could potentially be observed.

We found that the properties of perisaccadic perceptual mislocalization were modified if the appearance of the saccade target was altered. Moreover, this effect was not explained by different visibility of the perisaccadic high contrasts probe flashes as a function of the saccade-target appearance. We believe that these results motivate revisiting earlier neurophysiological investigations of corollary discharge from the SC (24, 25, 28, 58), but now from the perspective of saccades to images. For example, one could start mapping neurons in the pulvinar or medio-dorsal thalamus, which are involved in corollary discharge signaling from the SC to the cortex, but specifically during active vision tasks involving saccades to images rather than to spots. This could clarify if and how these brain areas may relay information about the visual appearance of saccade targets to their recipient neurons. It could also help explain what other roles these relay areas might have. For example, if SC visual responses, and not just saccade-related motor bursts, are relayed (24), then what is the purpose of such relaying, and what are the feature tuning properties of visual responses in areas like the pulvinar and the medio-dorsal thalamus?

While we did not focus on perisaccadic perceptual suppression too much in our current study, our second experiment already allowed us to measure perceptual suppression in time t1a (Figs. 6, 7). Interestingly, both the sample subject (Fig. 7A) and the population of subjects (Fig. 7B) showed that there was potentially stronger suppression (higher thresholds in the psychometric curves) when the saccade target had a low spatial frequency than when it had a high spatial frequency. This might suggest that even saccadic suppression strength itself, and not just perisaccadic perceptual mislocalization, can depend on the visual appearance of the saccade targets, which is consistent with our general hypotheses above, and also consistent with our observations with perisaccadic perceptual mislocalization in our main experiment. Indeed, we previously measured perceptual saccadic suppression with saccades across different visual textures, and we again observed stronger perisaccadic perceptual suppression with low spatial frequency textures (14). While our interpretations in that earlier study focused on the visual components of perisaccadic perceptual suppression, results from the current study and the sensory-tuned SC motor bursts (37) hint that motor-related components could also potentially account for at least some of these differences. Nonetheless, what is clear from our current perceptual detection experiments is that they cannot fully explain the mislocalization ones. This is because the probe contrasts in the mislocalization experiments were very high compared to when there could be any potential image dependencies of perceptual detection performance. In fact, at time t2, the perceptual detection psychometric curves were completely overlapping for the two different saccade-target image appearances (and almost fully recovered), but the perceptual mislocalization strength was still different.

Having said that, in our current experiments, the saccade target itself (and not just the representations of the probe flashes) was swept on the retina across saccades (from the periphery to the fovea in the case of the saccade target image). As a result, there was a motion sweep on the retina, which could potentially contribute to our observed perceptual results. In other words, if the motion sweep had different properties for the two different saccade target images, then it could still be possible that visual-visual interactions could account for our observations of different perceptual mislocalization across the image types. However, our gratings were orthogonal to the saccade vector direction. Thus, there was maximal retinal blurring of these gratings by the saccades. So, it seems less likely that this could have fully explained our mislocalization results. Moreover, our detectability experiments suggested that the probe contrast was so much above threshold contrast, especially at time t2, to be affected by potential subtle differences in the retinal motion sweeps associated with low and high spatial frequency grating saccade targets. Finally, the motion sweeps themselves might be independent of the mislocalization phenomenon in terms of mechanisms. For example, one could get mislocalization even with saccades towards a blank (61).

In any case, more generally, we believe that our experiments motivate the study of visual perception from an active perspective. Moreover, taking a more ecological approach to active vision than with simple dot stimuli would additionally be useful. It can reveal visual-visual and visual-motor interactions that are most relevant for natural behavior. Such an ecological approach could also include visual field asymmetries. For example, it is believed that over-representation of the upper visual field by the SC (21) could be ecologically relevant for active orienting across species (59, 62). Intriguingly, this motivated us to explore the effects of upper and lower visual field flashes in terms of mislocalization strength (Fig. 5), and we indeed found stronger perisaccadic perceptual mislocalization in the upper visual field. This is reminiscent of a differential effect also in saccadic suppression strength between the upper and lower visual fields (48), and it would be interesting in the future to further merge the topics of visual field asymmetries and perisaccadic vision but from a much more neurophysiological perspective.

## Acknowledgements

We were funded by the Deutsche Forschungsgemeinschaft (DFG; German Research Foundation) through the SFB 1233 Robust Vision (project number: 276693517). We were also funded by the DFG through the SPP 2205 Evolutionary Optimization of Neuronal Processing (project number: HA 6749/3-2) and the SPP 2411 Sensing LOOPS: Cortico-subcortical Interactions for Adaptive Sensing (project number: HA 6749/11-1).

